## Supplementary Material for "Loss of cell-autonomously secreted laminin-α2 drives muscle stem cell dysfunction in LAMA2-related muscular dystrophy"

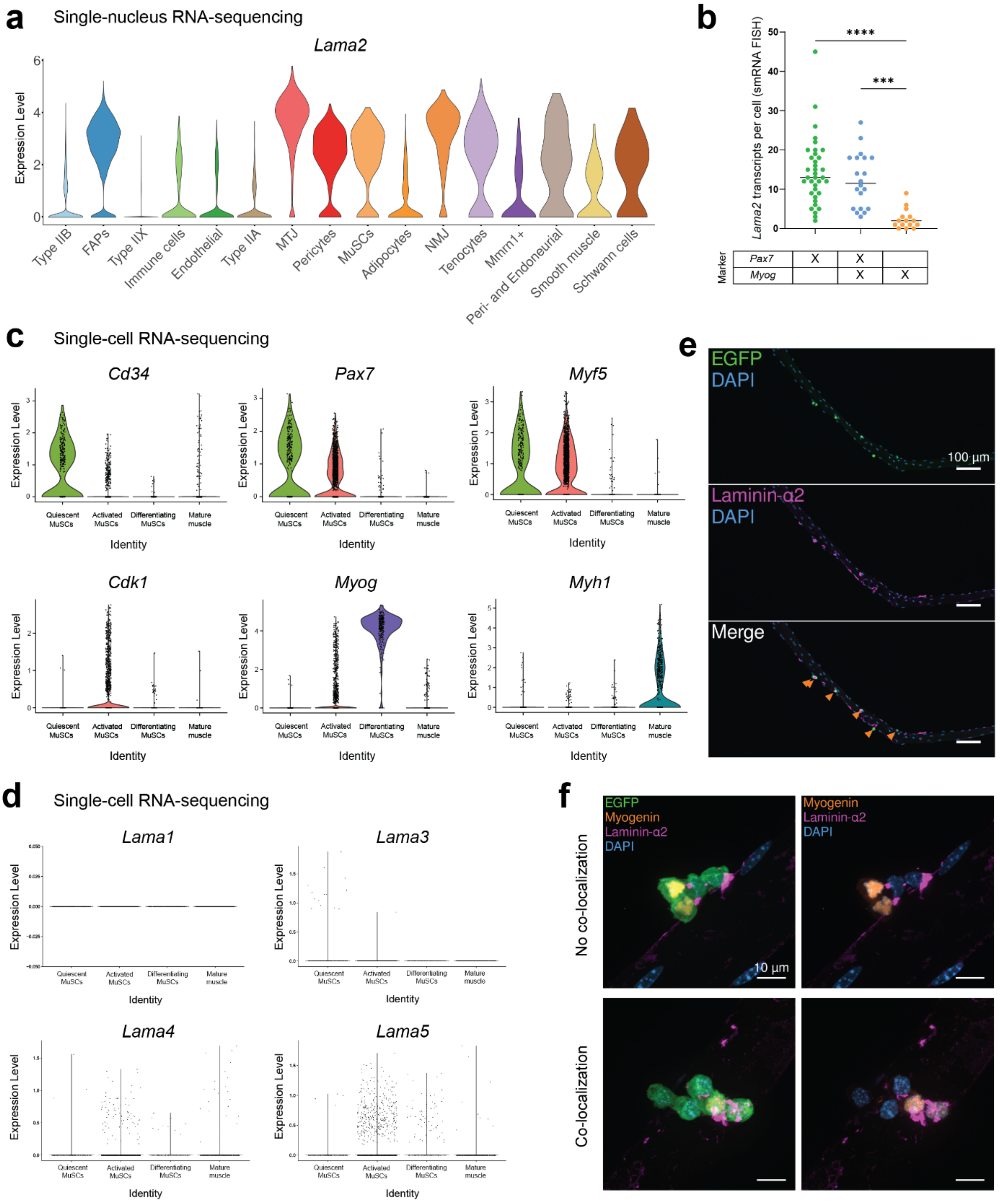

a
Single-nucleus RNA-sequencing
Lama2
Expression Level
Type IIB
FAPs
Type IIX
Immune cells
Endothelial
Type IIA
MTJ
Pericytes
MuSCs
Adipocytes
NMJ
Tenocytes
Mmrn1+
Peri- and Endoneurial
Smooth muscle
Schwann cells
b
Lama2 transcripts per cell (smRNA FISH)
****
***
Marker
Pax7
Myog
X
c
Single-cell RNA-sequencing
Cd34
Pax7
Myf5
Cdk1
Myog
Myh1
Quiescent MuSCs
Activated MuSCs
Differentiating MuSCs
Mature muscle
Identity
d
Single-cell RNA-sequencing
Lama1
Lama3
Lama4
Lama5
e
EGFP
DAPI
100 µm
Laminin-α2
DAPI
Merge
f
EGFP
Myogenin
Laminin-α2
DAPI
10 µm
No co-localization
Co-localization

**Fig. S1: Activated MuSCs express *Lama2* and remodel their microenvironment with laminin-α2.** **a** Violin plot showing *Lama2* expression in previously-defined cell types[1] of adult C57BL/6 mice. **b** Quantification of the number of *Lama2* mRNA transcripts detected by smRNA FISH in *Pax7*+/*Myog*-, *Pax7*+/*Myog*+ and *Pax7*-/*Myog*+ primary myoblasts *ex vivo*. Each dot represents one cell; cells were isolated from N = 2 mice. **c** Violin plots showing the expression of markers used to sub-cluster MuSCs in a published single-cell RNA-sequencing dataset[2]. **d** Violin plots showing the expression of *Lama* isoforms in the single-cell RNA-sequencing dataset[2]. **e** Immunostaining of an EDL fiber with EGFP-labelled MuSCs after 42 h in culture (EGFP in green, laminin-α2 in magenta, DAPI in blue). Orange arrows in the merge panel indicate EGFP+ MuSCs. **f** Immunostainings of EDL fibers with EGFP-labelled MuSCs at T72 showing no co-localization (1st row) and co-localization (2nd row) of Myogenin+/EGFP+ cells with laminin-α2 (EGFP in green, Myogenin in orange, laminin-α2 in magenta, DAPI in blue). In **b**, statistical significance was determined by one-way ANOVA with Tukey's multiple comparisons test. ***$P < 0.001$; ****$P < 0.0001$.

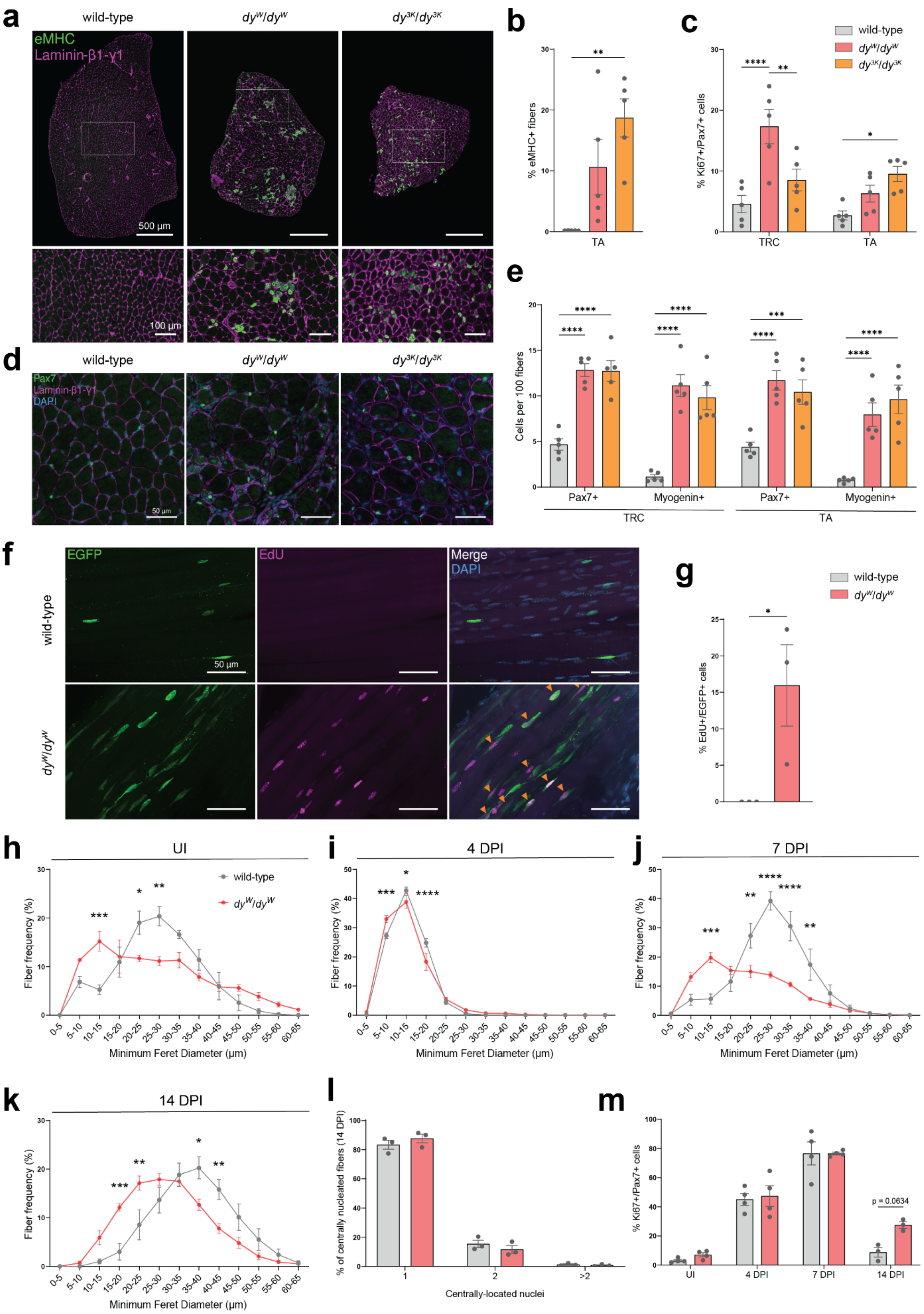

a
wild-type
$dy^W/dy^W$
$dy^{3K}/dy^{3K}$
eMHC
Laminin-β1-γ1
500 μm
100 μm
b
% eMHC+ fibers
TA
c
wild-type
$dy^W/dy^W$
$dy^{3K}/dy^{3K}$
% Ki67+/Pax7+ cells
TRC
TA
d
wild-type
$dy^W/dy^W$
$dy^{3K}/dy^{3K}$
Pax7
Laminin-β1-γ1
DAPI
50 μm
e
Cells per 100 fibers
Pax7+
Myogenin+
Pax7+
Myogenin+
TRC
TA
f
EGFP
EdU
Merge
DAPI
wild-type
$dy^W/dy^W$
50 μm
g
wild-type
$dy^W/dy^W$
% EdU+/EGFP+ cells
h
UI
wild-type
$dy^W/dy^W$
Fiber frequency (%)
Minimum Feret Diameter (μm)
i
4 DPI
Fiber frequency (%)
Minimum Feret Diameter (μm)
j
7 DPI
Fiber frequency (%)
Minimum Feret Diameter (μm)
k
14 DPI
Fiber frequency (%)
Minimum Feret Diameter (μm)
l
% of centrally nucleated fibers (14 DPI)
Centrally-located nuclei
1
2
>2
m
% Ki67+/Pax7+ cells
p = 0.0634
UI
4 DPI
7 DPI
14 DPI

**Fig. S2: LAMA2 MD mouse models have high levels of endogenous muscle repair. a** Representative immunostaining of regenerated muscle fibers in TA cross-sections of wild-type, $dy^{W}/dy^{W}$ and $dy^{3K}/dy^{3K}$ mice at 4-weeks-of-age (embryonic myosin heavy chain (eMHC) in green, laminin-β1-γ1 in magenta). **b** Quantification of the proportion of eMHC+ fibers at 4-weeks-of-age. **c** Quantification of the proportion of Ki67+/Pax7+ cells at 4-weeks-of-age. **d** Representative immunostaining of Pax7+ MuSCs in the TRC at 4-weeks-of-age (Pax7 in green, laminin-β1-γ1 in magenta, DAPI in blue). **e** Quantification of the number of Pax7+ and Myogenin+ cells per 100 fibers in the TRC and TA. **f** Wholemount immunostaining of EGFP-labelled MuSCs in the EDL after a 24 h EdU chase (EGFP in green, EdU in magenta, DAPI in blue). EdU+/EGFP+ cells are indicated by orange arrows in the merge panel. **g** Quantification of EdU+/EGFP+ MuSCs in EDL wholemounts. **h** – **k** Fiber size distribution in uninjured (UI) conditions (**h**), and at four (**i**), seven (**j**) and fourteen (**k**) days post-injury (DPI). **l** Quantification of the number of centrally-located nuclei in centrally-nucleated fibers at 14 DPI. **m** Quantification of the proportion of Ki67+/Pax7+ cells post-injury. Data are means ± SEM. In **b**, **c** and **e**, statistical significance was determined by one-way ANOVA with Tukey's multiple comparisons test. In **m**, statistical significance was determined by two-way ANOVAs with Bonferroni's multiple comparisons test. In **g**, **h**, **i**, **j**, **k** and **l**, statistical significance was determined by unpaired student's two-sided t-test. *$P < 0.05$; **$P < 0.01$; ***$P < 0.001$; ****$P < 0.0001$.

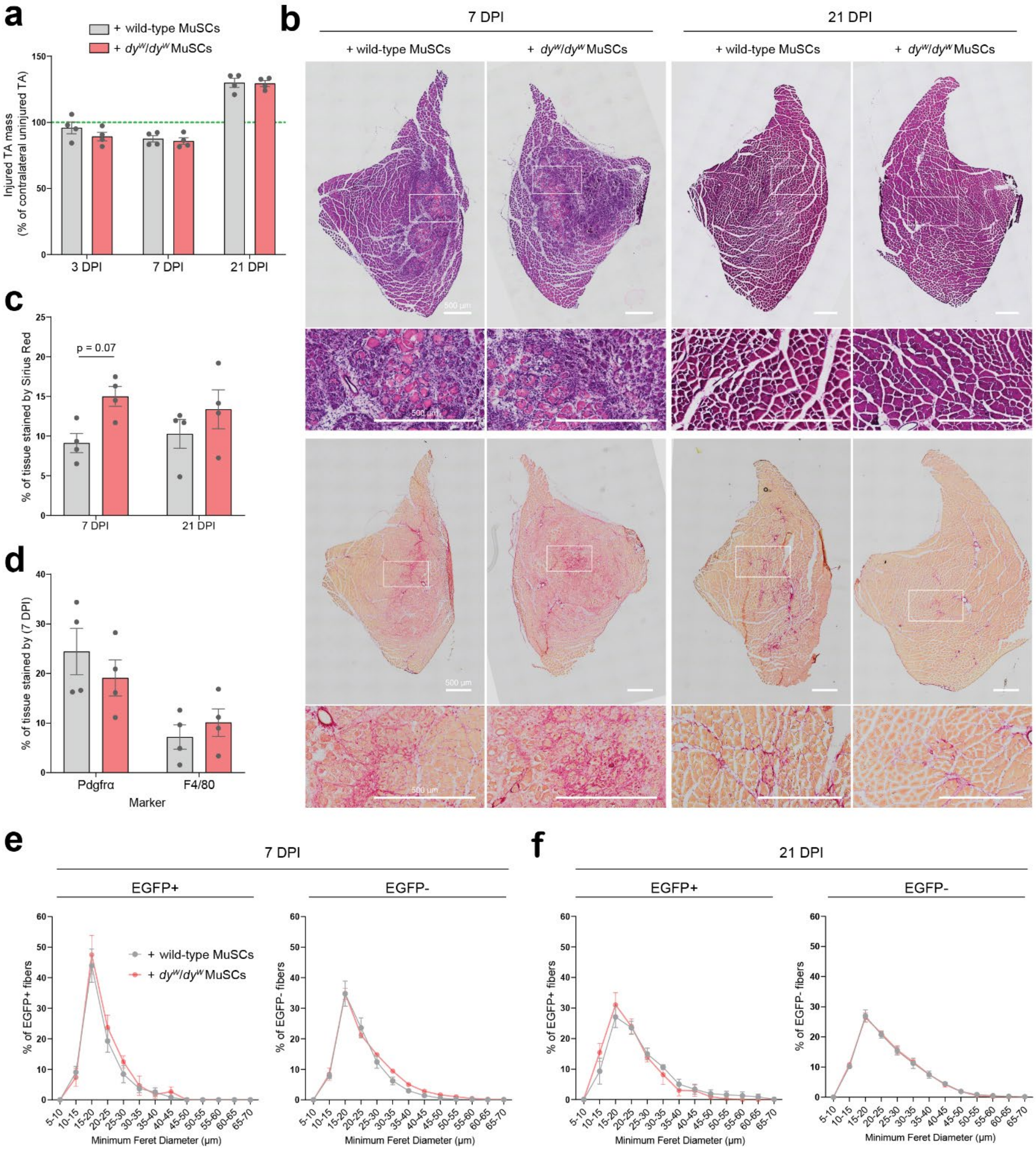
a
+ wild-type MuSCs
+ dyw/dyw MuSCs
Injured TA mass (% of contralateral uninjured TA)
3 DPI
7 DPI
21 DPI
b
7 DPI
21 DPI
+ wild-type MuSCs
+ dyw/dyw MuSCs
500 μm
c
% of tissue stained by Sirius Red
p = 0.07
d
% of tissue stained by (7 DPI)
Pdgfrα
F4/80
Marker
e
7 DPI
EGFP+
EGFP-
% of EGFP+ fibers
% of EGFP- fibers
Minimum Feret Diameter (μm)
f
21 DPI

**Fig. S3: Transplanted *$dy^W/dy^W$* MuSCs do not impair regeneration. a** Quantification of TA masses at 3, 7 and 21 days post-injury (DPI). Injured TA masses are shown as a percentage of the contralateral uninjured TA's mass (green dotted line). **b** Representative H&E (top) and Picro-Sirius Red (bottom) stains of recipient TA cross-sections at 7 (left) and 21 (right) DPI. **c** Quantification of the proportion of tissue stained by Sirius Red at 7 and 21 DPI in TA cross-sections. **d** Quantification of the proportion of tissue stained by Pdgfrα and F4/80 at 7 DPI in TA cross-sections. **e** Fiber size distribution of EGFP+ (left) and EGFP- (right) fibers at 7 DPI. **f** Fiber size distribution of EGFP+ (left) and EGFP- (right) fibers at 21 DPI. Data are means ± SEM. In **a** and **c**, statistical significance was determined by two-way ANOVAs with Bonferroni's multiple comparisons test. In **d**, **e** and **f**, statistical significance was determined by unpaired student's two-sided t-test.

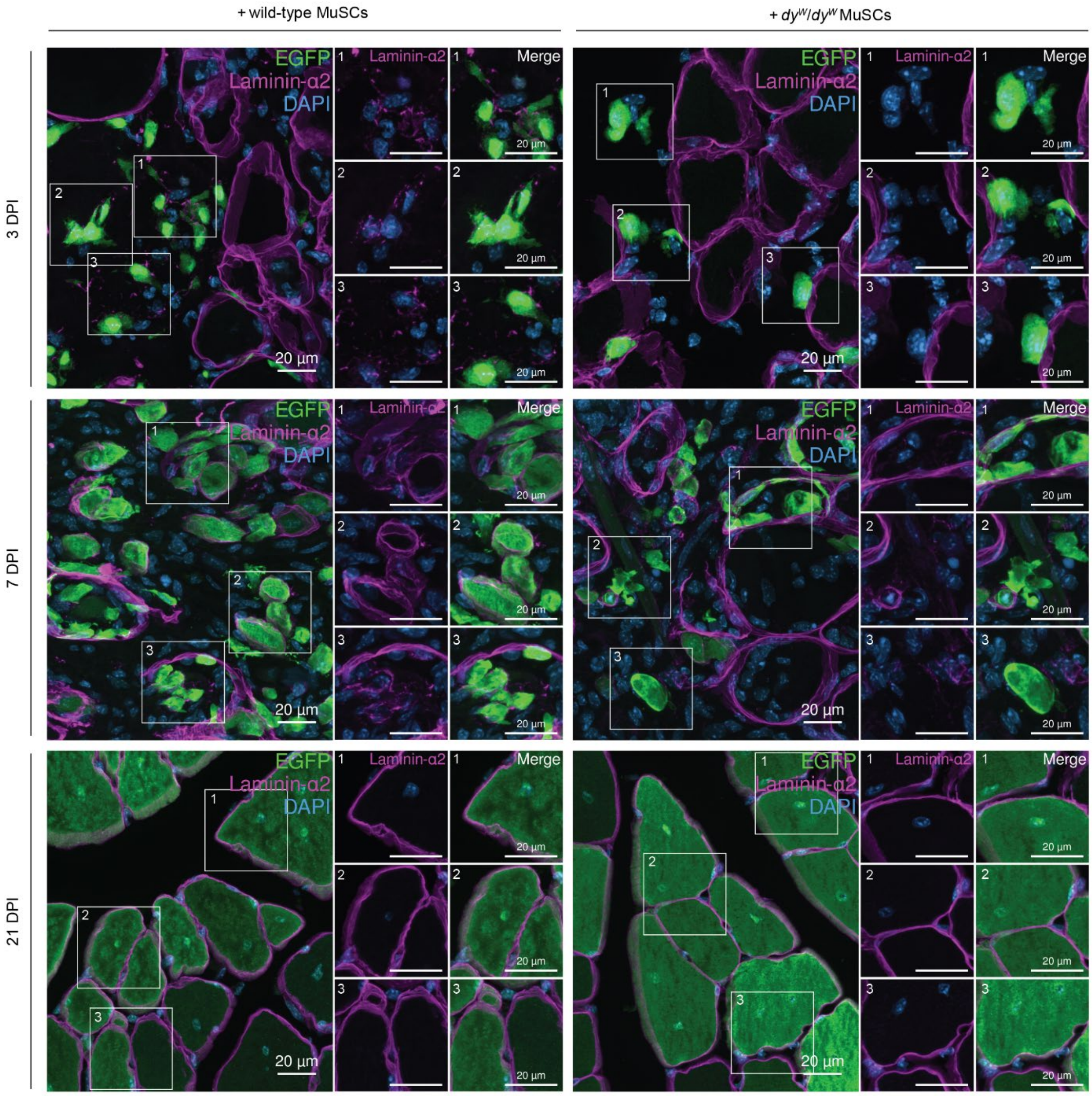


**Fig. S4: Proliferating wild-type MuSCs co-localize with laminin-α2 post-transplantation.** Representative immunostainings of recipient TA cross-sections at 3, 7 and 21 days post-injury (DPI) (EGFP in green, laminin-α2 in magenta, DAPI in blue). For each image, three zoomed-in panels are shown. At 3 DPI, laminin-α2 co-localizes with transplanted EGFP+ wild-type MuSCs, but not with transplanted EGFP+ *dy*$^W$/*dy*$^W$ MuSCs.

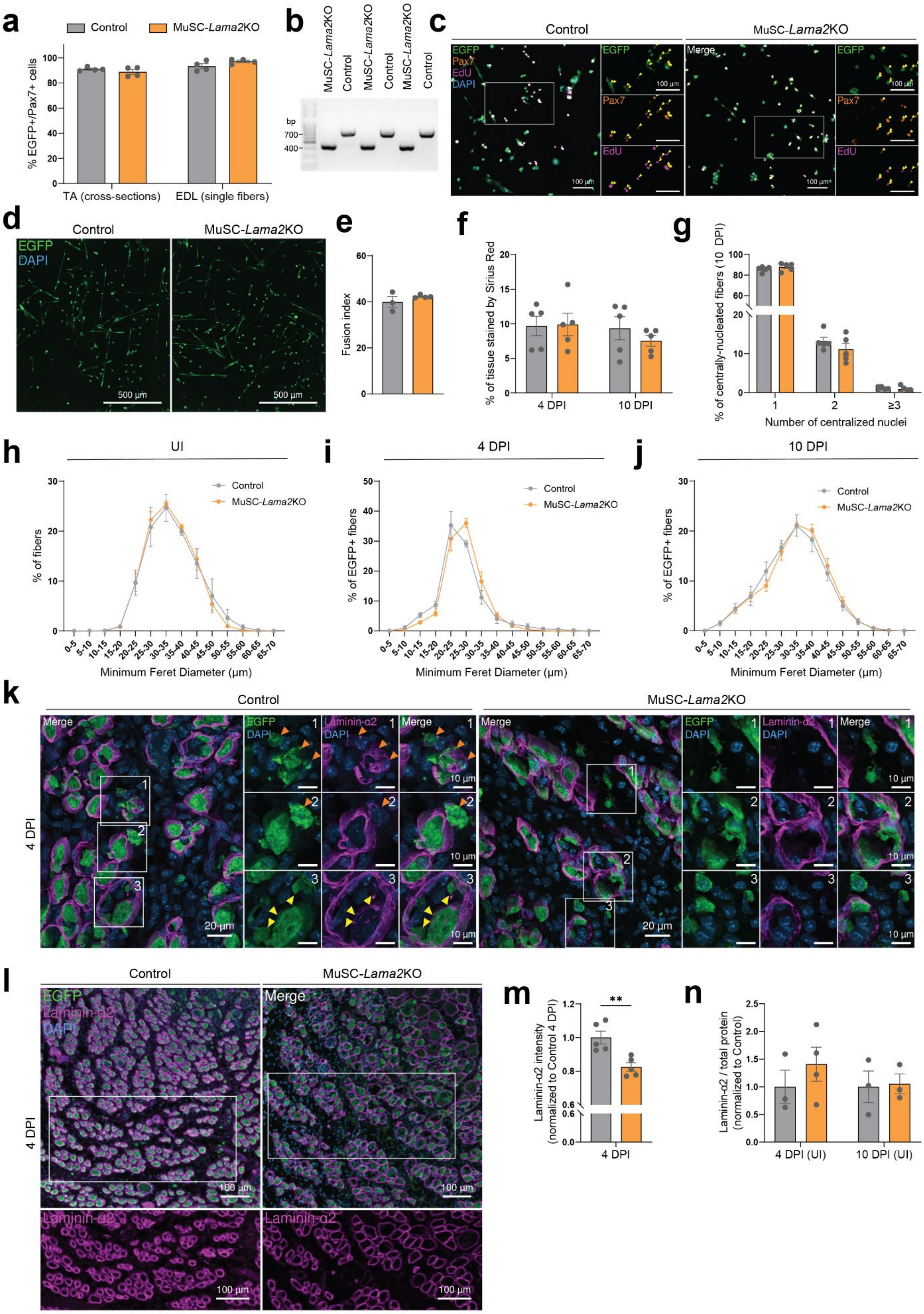

a
Control
MuSC-Lama2KO
% EGFP+/Pax7+ cells
TA (cross-sections)
EDL (single fibers)
b
MuSC-Lama2KO
Control
bp
700
400
c
Control
MuSC-Lama2KO
EGFP
Pax7
EdU
DAPI
Merge
100 µm
d
Control
MuSC-Lama2KO
EGFP
DAPI
500 µm
e
Fusion index
f
% of tissue stained by Sirius Red
4 DPI
10 DPI
g
% of centrally-nucleated fibers (10 DPI)
Number of centralized nuclei
≥3
h
UI
% of fibers
Minimum Feret Diameter (µm)
i
4 DPI
% of EGFP+ fibers
j
10 DPI
k
4 DPI
Laminin-α2
20 µm
10 µm
l
m
Laminin-α2 intensity (normalized to Control 4 DPI)
**
n
Laminin-α2 / total protein (normalized to Control)
4 DPI (UI)
10 DPI (UI)

**Fig. S5: MuSC-specific *Lama2* knockout is sufficient to impair MuSC proliferation and delay regeneration. a** Quantification of the proportion of cells that are double positive for Pax7 and EGFP in TA cross-sections and on single fibers isolated from the EDL after 5 consecutive days of tamoxifen treatment. **b** PCR on DNA isolated from sorted EGFP+ control and MuSC-*Lama2*KO cells demonstrating that *Lama2*'s exon 3 is specifically removed in MuSC-*Lama2*KO EGFP+ cells. Product size after deletion of exon 3 = 433 bp; product size of wild-type allele = 755 bp. **c** Representative immunostaining of cultured primary myoblasts after EdU incubation from 6-21 h post-plating (EGFP in green, Pax7 in orange, EdU in magenta, DAPI in blue). Yellow arrows indicate EdU+/Pax7+/EGFP+ cells. **d** Representative immunostaining of tissue culture dishes containing control and MuSC-*Lama2*KO cells after 3 days in differentiation medium. **e** Quantification of the fusion index (number of nuclei in myotubes divided by the total number of nuclei) after 3 days in differentiation medium. **f** Quantification of the proportion of tissue cross-section stained by Sirius Red at 4 and 10 days post-injury (DPI). **g** Quantification of the number of centrally-located nuclei at 10 DPI. **h – j** Fiber size distribution in uninjured (UI) conditions (**h**) and four (**i**) and ten days (**j**) post-cardiotoxin injury. **k** Representative immunostaining of TA cross-sections from control and MuSC-*Lama2*KO mice at 4 DPI. Orange arrows indicate laminin-α2 co-localization with interstitial EGFP+ cells; yellow arrows indicate laminin-α2 presence on the apical side of EGFP+ cells in ghost fibers. **l** Representative immunostaining showing reduced laminin-α2 signal in TA cross-sections of MuSC-*Lama2*KO mice 4 DPI (EGFP in green, laminin-α2 in magenta, DAPI in blue). **m** Quantification of laminin-α2 signal intensity in TA cross-sections at 4 DPI (average of 2 sections per mouse; N = 5 mice). **n** Quantification of laminin-α2 abundance in uninjured TAs of control and MuSC-*Lama2*KO mice at 4 and 10 DPI by Western blot (normalized to total protein and subsequently normalized to controls). Data are means ± SEM. In **a**, **e**, **g**, **h**, **i**, **j** and **m**, statistical significance was determined by unpaired student's two-sided t-test. In **f** and **n**, statistical significance was determined by two-way ANOVAs with Bonferroni's multiple comparisons test. ***P* < 0.01.

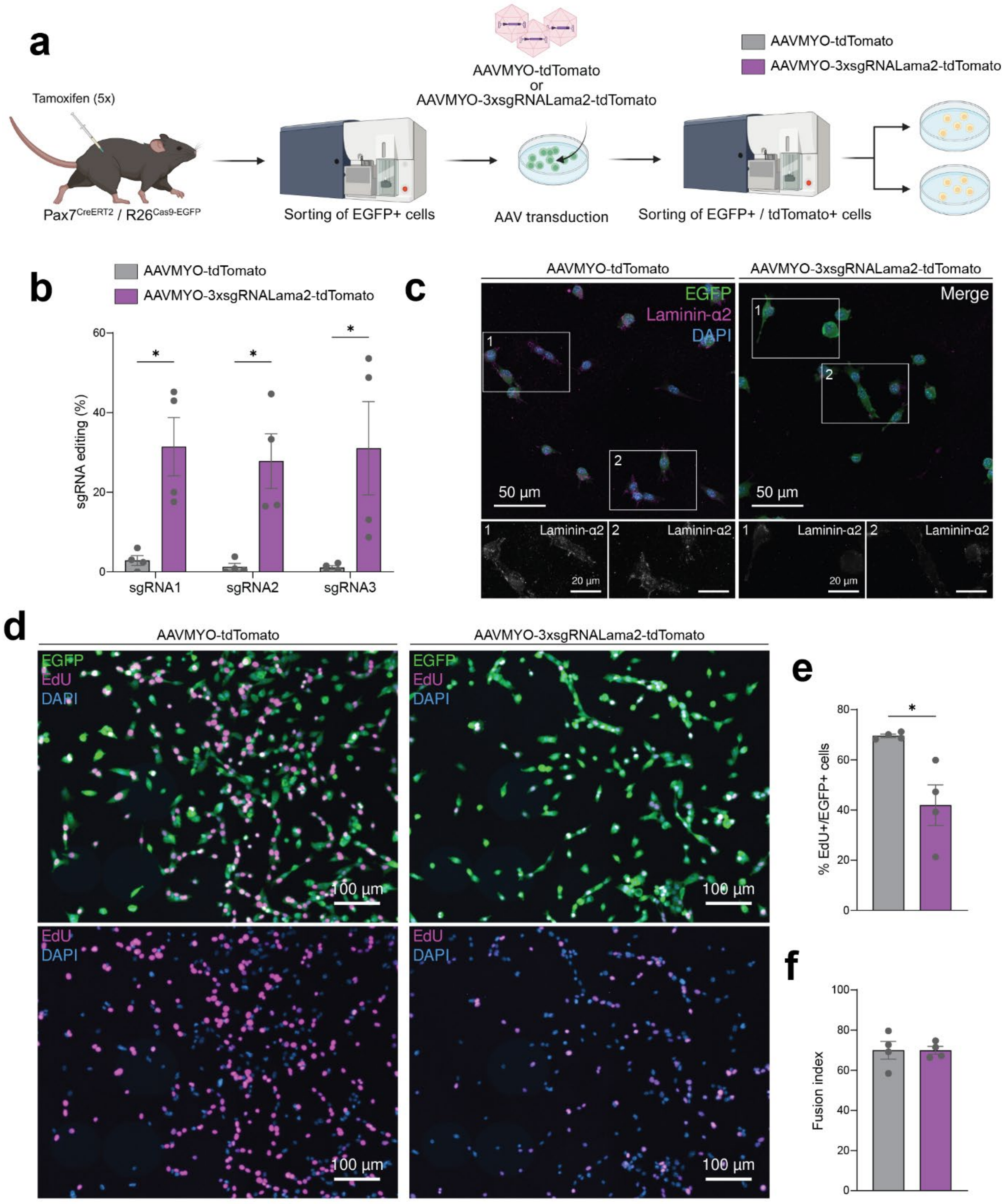
a
Tamoxifen (5x)
Pax7 CreERT2 / R26 Cas9-EGFP
Sorting of EGFP+ cells
AAVMYO-tdTomato
or
AAVMYO-3xsgRNALama2-tdTomato
AAV transduction
Sorting of EGFP+ / tdTomato+ cells
AAVMYO-tdTomato
AAVMYO-3xsgRNALama2-tdTomato
b
AAVMYO-tdTomato
AAVMYO-3xsgRNALama2-tdTomato
sgRNA editing (%)
sgRNA1
sgRNA2
sgRNA3
c
AAVMYO-tdTomato
AAVMYO-3xsgRNALama2-tdTomato
EGFP
Laminin-α2
DAPI
Merge
50 µm
Laminin-α2
20 µm
d
AAVMYO-tdTomato
AAVMYO-3xsgRNALama2-tdTomato
EGFP
EdU
DAPI
100 µm
e
% EdU+/EGFP+ cells
f
Fusion index

**Fig. S6: CRISPR/Cas9-mediated *Lama2* knockout reduces MuSC proliferation *ex vivo*.**
**a** Experimental approach: mice with inducible MuSC-specific expression of Cas9 and EGFP[3, 4] were treated with tamoxifen for 5 consecutive days. EGFP+ cells were then isolated *via* FACS, expanded on collagen-coated tissue culture dishes before AAVMYO was added to medium to deliver either a CMV-driven tdTomato transgene, or a CMV-driven tdTomato transgene and the sequence for 3 single guide RNAs (sgRNAs) targeting exons 2 and 3 of *Lama2*. Four days after the addition of AAVMYO, the cells were passaged. Transduced EGFP+/tdTomato+ cells were sorted *via* FACS and re-plated to assess the effect of *Lama2* mutations on isogenic PMs *ex vivo*. **b** Quantification of sgRNA editing efficiency by Tracking of Indels by Decomposition (TIDE) (see Methods for more details). **c** Representative immunostaining of laminin-α2 in tissue culture dishes containing isogenic EGFP+ cells that received either AAVMYO-tdTomato or AAVMYO-3xsgRNALama2-tdTomato (EGFP in green, laminin-α2 in magenta, DAPI in blue). Boxes 1 and 2 are zoomed-in panels showing laminin-α2 in white. **d** Representative immunostaining of isogenic EGFP+ cells that received either AAVMYO-tdTomato or AAVMYO-3xsgRNALama2-tdTomato. Cells were fixed after a 15 h incubation with EdU (EGFP in green, EdU in magenta, DAPI in blue). **e** Quantification of the proportion of EdU+/EGFP+ cells after a 15 h incubation with EdU. **f** Quantification of the fusion index (number of nuclei in myotubes divided by the total number of nuclei) after 7 days of differentiation. Data are means ± SEM. In all graphs, statistical significance was determined by unpaired student's two-sided t-test. *$P < 0.05$. Experimental scheme **a** was created with Biorender.com.

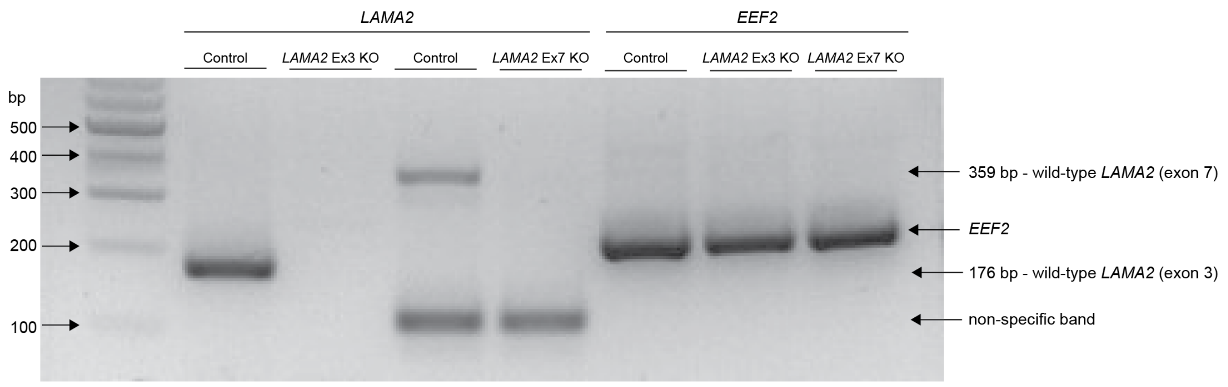


**Fig. S7: Loss of *LAMA2* mRNA in *LAMA2* knockout human induced pluripotent stem cells.** RT-PCR showing a loss of *LAMA2* mRNA in *LAMA2* Ex3 KO and *LAMA2* Ex7 KO human induced pluripotent stem cells. The expression of *EEF2* is shown for all samples as a control. In the comparison of *LAMA2* expression in control versus *LAMA2* Ex3 KO cells, the forward primer targets exon 2 and the reverse primer targets exon 3 (product length = 176 bp). In the comparison of *LAMA2* expression in control versus *LAMA2* Ex7 KO cells, the forward primer targets exon 6 and the reverse primer targets exon 8 (product length =359 bp; the lower band detected in both samples is non-specific and was detected in two repeats of the experiment). For the comparison of *EEF2* expression in all samples product length is 173 bp.

**Supplementary Table 1: List of antibodies used.**

| Antibody | Company | Reference | Dilution (Immunofluorescence) | Dilution (Western blot) | Dilution (FACS) |
|---|---|---|---|---|---|
| Mouse anti-Pax7 | DSHB | PAX7 | 1/20 | N/A | N/A |
| Rabbit anti-Laminin β1-γ1 | Sigma-Aldrich | L9393 | 1/200 | N/A | N/A |
| Mouse anti-embryonic myosin heavy chain | DSHB | F1.652 | 1/100 | N/A | N/A |
| Rabbit anti-Ki67 | abcam | ab15580 | 1/100 | N/A | N/A |
| Rabbit anti-Myogenin | abcam | ab12480 | 1/400 | N/A | N/A |
| Chicken anti-GFP | Molecular Probes | A10262 | 1/400 | N/A | N/A |
| Anti-Laminin-2 (α-2 Chain) antibody, Rat monoclonal | Sigma | L0663 | 1/200 | N/A | N/A |
| Laminin alpha-2 Monoclonal Antibody (CL3450) | Invitrogen | MA5-24656 | N/A | 1/1,000 | N/A |
| Rabbit anti-Desmin | abcam | ab15200 | 1/400 | N/A | N/A |
| Rabbit anti-Human/Mouse Cleaved Caspase-3 (Asp175) | R&D Systems | MAB835 | 1/100 | N/A | N/A |
| Rat anti F4/80 (Cl:A3-1) | abcam | ab6640 | 1/100 | N/A | N/A |
| MF20 | DSHB | MF 20 | 1/100 | N/A | N/A |
| Cy5 AffiniPure Goat anti-Rabbit IgG (H+L) | Jackson | 111-175-144 | 1/500 | N/A | N/A |
| Cy3 AffiniPure Goat anti-Mouse IgG1 | Jackson | 115-165-205 | 1/500 | N/A | N/A |
| Cy3 AffiniPure Goat anti-Mouse IgG (H+L) | Jackson | 115-165-003 | 1/500 | N/A | N/A |
| Cy3 AffiniPure Goat anti-Rat IgG (H+L) | Jackson | 112-165-143 | 1/500 | N/A | N/A |
| AF488 AffiniPure Goat anti-Mouse IgG1 | Jackson | 115-545-205 | 1/500 | N/A | N/A |
| AF488 AffiniPure Goat anti-Chicken IgG (H+L) | Jackson | 103-545-155 | 1/500 | N/A | N/A |
| Peroxidase AffiniPure™ Goat Anti-Mouse IgG, light chain specific | Jackson | 115-035-174 | N/A | 1/10,000 | N/A |
| Rat anti-Integrin α7, AF647 conjugated | ablab | 67-0010-05 | N/A | N/A | 1/1000 |
| Rat anti-Ly-6A/E, PE conjugated | BD Pharmingen | 553336 | N/A | N/A | 1/500 |
| Mouse anti-CD31, PE conjugated | BD Pharmingen | 555027 | N/A | N/A | 1/500 |
| Rat anti-CD45, PE conjugated | eBioscience | 12-0451-82 | N/A | N/A | 1/500 |
| Rat anti-CD11b, PE conjugated | eBioscience | 12-0112-81 | N/A | N/A | 1/500 |

**Supplementary Table 2: List of primers used.**

| Gene | Figure | Forward | Reverse |
|---|---|---|---|
| *Lama2* | 1b | TGCCCTTTCTCACCCACCCTT | GTTGATGCGCTTGGGAC |
| *Pax7* | 1b | GAGGTGACAGGAGGCAGAAG | AGCTGCCAGCAAGATGGTAT |
| *Myog* | 1b | ACTCCCTTACGTCCATCGTG | CAGGACAGCCCCACTTAAAA |
| *Gapdh* | 1b | ACCCAGAAGACTGTGGATGG | GGATGCAGGGATGATGTTCT |
| *Lama2 exon 3* | S5b | AGCACCTTTCCAACAGGAGA | TGTGCTGTTGTGTTCCCTTC |
| TIDE sgRNA 1+2 | S6b | AAGGCTGGTGGTCAGTGTTT | TCAGCATCGCTCCCAACTTT |
| TIDE sgRNA 3 | S6b | AGGACCCGAGATGTACTGCA | TCTGTGGCCAGGGAGTCTAA |
| *LAMA2* (control vs *LAMA2* Ex 3 KO) | S7 | CGACCAATGCAACATGTGGAG | CTGCCACCAAGTGTTCTTTCCA |
| *LAMA2* (control vs *LAMA2* Ex 7 KO) | S7 | TTTCAGTTGGAGGGATGTGC | GGGTCTGAAGAAGCCATCAGT |
| *EEF2* | S7 | TCAGCACACTGGCATAGAGG | GACATCACCAAGGGTGTGC |

**Supplementary Table 3: List of guides used.**

| Use | Sequence |
|---|---|
| *Lama2* KO in murine MuSCs - guide 1 | CCGATTACGAATGCTATTGA |
| *Lama2* KO in murine MuSCs - guide 2 | CAGAGTCCCAGTATCAAGAA |
| *Lama2* KO in murine MuSCs - guide 3 | GATTGCAGATTCGGCACTGA |
| *LAMA2* Ex3 KO in hiPSCs – guide 1 | TCATCTTATCAAGAAAACAATGG |
| *LAMA2* Ex3 KO in hiPSCs – guide 2 | CTGTATGGTGCTATGAGACAAGG |
| *LAMA2* Ex7 KO in hiPSCs – guide 1 | CCTAGAGGCCTAGGAATCAAC |
| *LAMA2* Ex7 KO in hiPSCs – guide 2 | GACCAGTTGTCACTATTAGGCT |
